# Long-term euxinia hinders microbial ammonium removal in brackish coastal waters

**DOI:** 10.1101/2025.06.30.662269

**Authors:** Nicky Dotsios, Lina Piso, Jessica Venetz, Olga M. Żygadłowska, Niels A.G.M. van Helmond, Wytze K. Lenstra, Claudia Frey, Alessandra Mazzoli, Moritz F. Lehmann, Florian Roth, Christian Stranne, Christoph Humborg, Tom Berben, Pedro E. Leão, Maartje A.H.J. van Kessel, Sebastian Lücker, Mike S.M. Jetten, Caroline P. Slomp

**Author notes:** These authors contributed equally to this work (Nicky Dotsios, Lina Piso).

## Abstract

Anthropogenic activities are key drivers of eutrophication and deoxygenation in coastal marine ecosystems. This stimulates the anaerobic degradation of organic matter and the release of reduced products, such as ammonium, methane, and hydrogen sulfide, which may, in turn, exacerbate eutrophication and deoxygenation. In this study, using a combination of chemical and microbial analyses, we assess the nitrogen dynamics in the water column of a eutrophic coastal system (Stockholm Archipelago) at three sites with contrasting redox conditions (oxic to long-term euxinic). At the oxic site, counter gradients of ammonium and oxygen in the water column, low nitrate δ^15^N values in bottom waters, and the 16S rRNA gene-based presence of nitrifiers indicate nitrification near the sediment-water interface. At the seasonally and long-term euxinic sites, nitrification, as inferred from the water column oxygen and nutrient profiles and the relative abundance of nitrifiers, primarily occurred near the oxycline. At these two sites, nitrate was removed below the oxycline through denitrification linked to sulfide oxidation by *Sulfurimonas*. Nitrous oxide emissions from surface waters in the archipelago reached up to 40 µmol m^-2^ d^-1^ and were not directly related to water column redox conditions, indicating that multiple factors control coastal emissions of this greenhouse gas to the atmosphere.

The relative abundance of 16S rRNA genes and of N-cycle genes in metagenomes was highest at the seasonally euxinic site. Importantly, nitrifiers were significantly less abundant at the long-term euxinic site. Our results highlight that prolonged euxinia promotes recycling of ammonium over its removal, likely due to sulfide inhibition of nitrification, which sustains eutrophication and deoxygenation of coastal systems.

## Introduction

Coastal ecosystems worldwide are increasingly suffering from human-induced eutrophication (Breitburg et al., 2018). Eutrophication refers to an over-enrichment of nutrients, primarily nitrogen and phosphorus, from sources such as agricultural runoff and sewage discharge (Breitburg et al., 2018; Rabalais et al., 2009). Coastal eutrophication forms a threat because algal growth is stimulated and, upon decay of the biomass, dissolved oxygen (O_2_) is consumed, leading to hypoxic (O_2_ < 63 µM), anoxic (O_2_ = 0 µM), or euxinic conditions (presence of hydrogen sulfide [H_2_S]) (Rabalais et al., 2009, 2010). Deoxygenation can have severe consequences for marine life, such as habitat loss and alterations in food web structure (Diaz & Rosenberg, 2008; Rabalais et al., 2009).

Coastal eutrophication and deoxygenation stimulate the release of ammonium (NH_4_^+^), methane (CH_4_), and H_2_S from sediments (Conley et al., 2007). This contributes to the degradation of coastal ecosystems because recycled NH_4+_ can promote further algal blooms, while CH_4_ is a potent greenhouse gas, and H_2_S is highly toxic. Moreover, the oxidation of these reduced compounds can stimulate additional O_2_ loss (Hermans et al., 2019). Given the role of NH_4_^+^ as both a nutrient and reductant, it is essential to understand its dynamics in coastal systems, along with those of the other key components of the nitrogen cycle, all of which depend on ammonia oxidation as the first step.

Nitrogen cycling in coastal systems involves a range of microbial pathways, including aerobic ammonia oxidation (or nitrification), and anaerobic pathways, such as denitrification, dissimilatory nitrate reduction to ammonium (DNRA), and anaerobic ammonium oxidation (anammox). Ammonia-oxidizing bacteria (AOB) or archaea (AOA) oxidize ammonia to nitrite, with *amo*A and *hao* as diagnostic genes encoding for subunits of the key proteins ammonia monooxygenase and hydroxylamine dehydrogenase, respectively. Nitrite-oxidizing bacteria (NOB) subsequently oxidize this nitrite to nitrate using nitrite oxidoreductase (NxrABC) as a key protein. While nitrification is an aerobic process, nitrifying microorganisms have the ability to adapt, survive, and function under low-oxygen conditions (Jäntti et al., 2018). In marine environments, AOA, mainly comprising *Candidatus* Nitrosopumilus maritimus, often outcompete AOB (Labrenz et al., 2010; Santoro et al., 2010). However, canonical AOB of the genus *Nitrosomonas* may be favoured in freshwater and brackish waters with relatively high NH_4_^+^ concentrations (De Bie et al., 2001; Freitag et al., 2005).

Denitrification is a multi-step process that reduces nitrate (NO_3_^-^) to dinitrogen gas (N2) via nitrite (NO_2_^-^), nitric oxide (NO), and nitrous oxide (N_2_O). The separate steps of denitrification are catalyzed by, respectively, nitrate reductase (encoded by *narG* or *napAB*), nitrite reductase (*nirK* or *nirS*), NO reductase (*norBC* [cNOR] or *qnorB*/*norZ* [qNOR]), and nitrous oxide reductase (*nos*Z). During DNRA, NO_2_^-^ is directly converted to ammonia by a multiheme cytochrome c protein complex encoded by a *nrfAH*. Anammox, on the other hand, directly converts NH_4_^+^ and NO_2_^-^ to N_2_, with NO and hydrazine (N_2_H_4_) as important intermediates (Kartal et al., 2011). Anammox is carried out by bacteria that thrive in oxygen-depleted environments, such as stratified coastal water bodies, oxygen minimum zones (OMZs), and anoxic sediments (Dalsgaard et al., 2012; Jensen et al., 2008; Kuypers et al., 2003).

Accumulation of H_2_S in coastal systems can inhibit a wide range of microorganisms, including AOB, NOB, and anammox bacteria (Jensen et al., 2008; Lam et al., 2007; Russ et al., 2019). Denitrifying bacteria and DNRA microorganisms are more tolerant towards H_2_S and may even couple H_2_S oxidation to nitrate reduction (Bonaglia et al., 2016; Brettar et al., 2006; Grote et al., 2007; Kox & Jetten, 2015; Russ et al., 2014). Importantly, nitrous oxide reductases (NosZ) are inhibited by H_2_S (Bartacek et al., 2010), potentially contributing to enhanced emissions of the greenhouse gas nitrous oxide from euxinic coastal waters to the atmosphere (Dalsgaard et al., 2013). Additionally, the inhibition of nitrification by H_2_S (Joye & Hollibaugh, 1995; Lam et al., 2007) may lead to increased recycling of nitrogen in coastal systems (Kemp & Testa, 2011). While many studies have focused on nitrogen cycling at redox interfaces in coastal seas (e.g., Dalsgaard et al., 2013; Frey et al., 2014; Hietanen et al., 2012; Jäntti et al., 2018), the dynamics of O_2_, H_2_S, and nitrogen in periodically stratified coastal waters, where periods of anoxia and H_2_S accumulation (euxinia) alternate with periods of oxygenation, are less well understood.

In this study, we investigate key microbial processes controlling nitrogen dynamics in the water column at three sites in the Stockholm Archipelago with contrasting water column redox conditions (oxic to long-term euxinic). We combine water column depth profiles of O_2_, H_2_S, NH_4_^+^, NO_2_^-^, NO_3_^-^, N_2_O, and δ^15^N and δ^18^O-NO_3_^-^ with microbial abundances determined through 16S rRNA gene and metagenomic sequencing to determine zones of potential N-transformation. Rates of N_2_O emission to the atmosphere are quantified from continuous *in situ* measurements of N_2_O in surface waters of the archipelago. Our findings highlight that euxinia acts as a key control on microbial nitrogen dynamics in the water column of this eutrophic coastal system, promoting nitrogen recycling over removal.

## Materials and Methods

### Study area

The Stockholm Archipelago is a brackish-water system, consisting of thousands of islands enclosed by a network of basins connected through straits (Almroth-Rosell et al., 2016; Wåhlström et al., 2024). The western part of the archipelago, which has a surface water salinity of ∼4.5 (Żygadłowska et al., 2024), receives freshwater from Lake Mälaren via the Norrström River (Almroth-Rosell et al., 2016) and brackish water from the Baltic Sea. High nutrient input from land has resulted in widespread eutrophication (Almroth-Rosell et al., 2016; Van Helmond et al., 2020). Water column stratification typically develops in summer and weakens in fall and winter (Gidhagen, 1987), leading to the development of hypoxia, anoxia, or euxinia below the thermocline in many parts of the archipelago (Conley et al., 2011). This was confirmed based on depth profiling at 11 stations (Fig. 1a; Żygadłowska et al., 2024).

**Figure 1.**
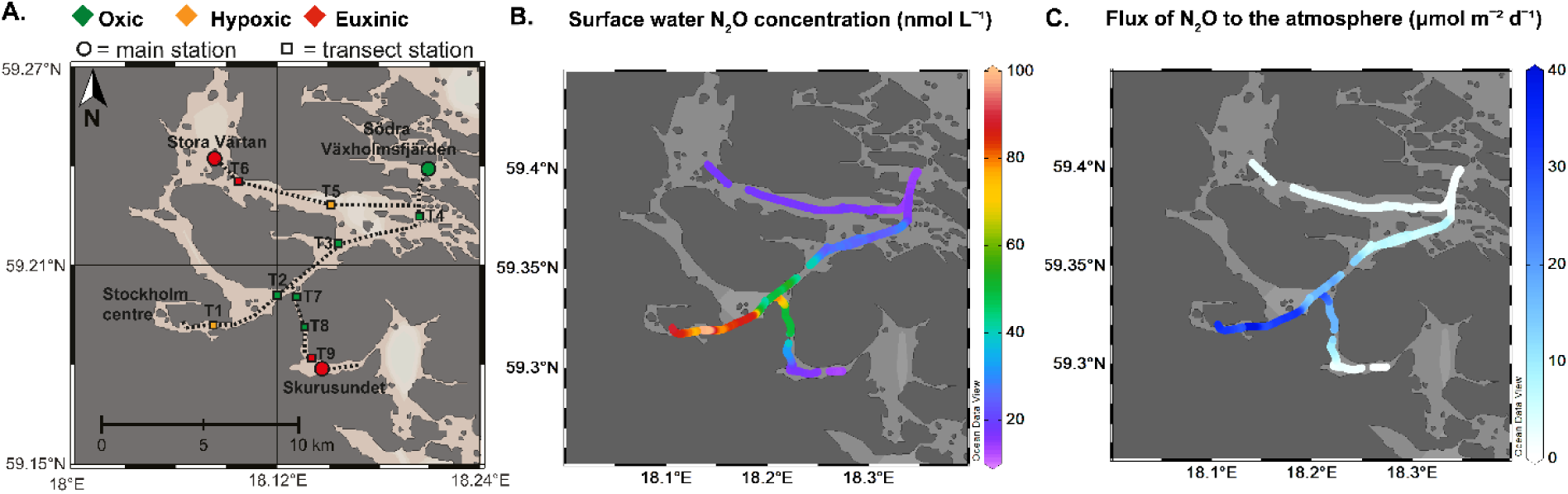
(a) Map of the study area in the Stockholm Archipelago (Żygadłowska et al 2024). Continuous N_2_O measurements were conducted along three transects (indicated with the dotted lines). The three study sites are indicated with circles, additional transect stations are indicated with squares. Bottom water redox conditions are indicated with a color scale ranging from oxic (green) to euxinic (red). (b) Surface water N_2_O concentrations from continuous measurements using the WEGA system. (c) Calculated N_2_O flux to the atmosphere based on the WEGA measurements.

In this study, continuous surface water N_2_O concentration measurements were performed along transects in the Stockholm Archipelago with R/V Electra in September 2022 (Fig. 1). Water column profiles of key solutes were determined at three sites (Table 1), selected based on data from long-term monitoring (SMHI, 2020; Żygadłowska et al., 2024).

**Table 1.** General characteristics of the three study sites in the Stockholm Archipelago at the time of sampling.

| Study site | Coordinates | Water column redox conditions | Water depth (m) |
| --- | --- | --- | --- |
| Södra Vaxholmsfjärden | 59.3995 °N<br>18.3472 °E | Oxic | 22 |
| Stora Värtan | 59.4043 °N<br>18.1367 °E | Seasonally euxinic | 27 |
| Skurusundet | 59.2985 °N<br>18.2295 °E | Long-term euxinic | 24 |

### Continuous surface water measurements and sea-air flux of N_2_O

The N_2_O concentration in the surface water was continuously measured along three transects in the archipelago (Fig. 1b) using a water equilibrator gas analyzer (WEGA) system (model G2131-I, Picarro Inc; Żygadłowska et al. 2024). The measured N_2_O concentrations were used to calculate diffusive fluxes of N_2_O to the atmosphere along the transect (Fig. 1c), as detailed in the supplementary information. The WEGA system was also used to measure depth profiles of N_2_O in the H_2_S-free part of the water column at the three sampling sites.

### Water column sampling

Depth profiles of temperature, salinity, and dissolved O_2_ were obtained with a CTD (Sea-Bird SBE 911 Plus, Sea-Bird Electronics, Bellvue, WA, USA) equipped with an O_2_ sensor (Seabird SBE 43, Sea-Bird Electronics). Water column samples were retrieved at a depth resolution of 1-2 m on the upcast of the CTD using a Rosette sampler with 12 Niskin bottles of 5 L. Samples for H_2_S were filtered through 0.2 μm nylon membrane filters into glass vials with zinc acetate solution (2%) and stored at 4 °C (Żygadłowska et al., 2024). Samples for NH_4_^+^, NO_2_^-^, and NO_3_^-^ were filtered through 0.2 μm nylon membrane filters into 50 mL polypropylene tubes and stored at −20 °C (Żygadłowska et al., 2023). Samples for dissolved Fe and Mn were filtered using a nylon membrane filter with a diameter of 0.2 μm into 250 ml acid-washed wide-mouth LDPE Bottles and acidified with 200 μL concentrated ultrapure HCl per 100 mL of sample (Żygadłowska et al., 2023). Samples for total Fe and Mn were collected in the same manner but without the filtration step. Samples for CH_4_ were collected in borosilicate serum bottles of 125 mL using norprene tubing, allowing them to overflow. The bottles were quickly closed with butyl rubber stoppers and aluminum crimp caps, poisoned with 0.25 mL saturated zinc chloride solution and stored upside down in the dark (Żygadłowska et al., 2024). Samples for DNA extractions were collected into 1L sterile plastic bottles and stored at 4 °C. Within 24 hours, between 0.5 and 1L of water was filtered on Supor® PES filters (pore size 0.22 µm; diameter 47 mm; Pall Lab) using a vacuum pump. The filters were flash frozen at −80 °C and stored at −20 °C until further processing.

### Chemical analysis of water samples

H_2_S concentrations were determined using the phenylenediamine and ferric chloride method (Cline, 1969). Concentrations of NH_4_^+^ were determined with the indophenol blue method (Solorzano, 1969). NO_3_^-^ and NO_2_^-^ concentrations were measured by two methods. In one approach, reduction to NO was followed by detection using a Nitric Oxide analyzer (NOA280i, GE Analytical Instruments, Manchester, UK). In the second method, a modified Griess assay was used, mixing a 1:1 ratio of reagents A and B with one volume of sample. Reagent A consists of 1% sulfanilic acid in 1 M HCL, and reagent B is a 0.1% (w/v) N-(1- naphtyl) ethylenediamine solution. After incubation for 10 minutes and the absorbance was measured at 540 nm on a SpectraMax190 plate reader (Molecular Devices, San Jose, US). To measure NO_3_^-^ in the same sample, NO_3_^-^ was reduced to NO_2_^-^ by adding 6.9 mM VCl_3_ and incubation for 30 min at 60 °C. Absorbance was then remeasured at 540 nm (Griess, 1879; Wang et al., 2016). Total and dissolved Fe and Mn concentrations were measured by inductively coupled plasma mass spectrometry (ICP-MS, Perkin Elmer NexION 2000). Particulate Fe and Mn were determined as the difference between total and dissolved metals (Żygadłowska et al., 2023).

### Isotopic analysis

For δ^15^N and δ^18^O-NO_3_^-^ analysis, the denitrifier method was used (Casciotti et al., 2002; Sigman et al., 2001). This method utilizes a denitrifying bacterial strain that lacks N_2_O- reductase activity (*Pseudomonas aureofaciens*; ATC 13985) to convert NO_3_^-^ and NO_2_^-^ to N_2_O. The generated N_2_O was analyzed using a continuous flow isotope ratio mass spectrometer (CF-IRMS, Delta V Plus, Thermo) using the procedure described by Frey et al. (2014). As the denitrifier method yields the combined isotopic composition of NO_2_^-^ and NO_3_^-^, the samples were first treated with sulfamic acid (10 μL of 8% sulfamic acid per mL of sample) to remove NO_2_^-^ and avoid offsets between the NO_3_^-^ concentration in the sample and the produced N_2_O (Casciotti & McIlvin, 2007). After NO_3_^-^ reduction, the sample volume was adjusted to contain 10 nmol N_2_O per sample (Frey et al., 2014). Only samples with an initial NO_3_^-^ concentration higher than 1 μmol L^-1^ were analysed with the IRMS to avoid interferences by other solutes and ensure the highest sensitivity of the analysis (Casciotti et al., 2002). Samples from the oxic site were below the detection limit of the assay. Each sample was measured in triplicate and corrected for the blank and for oxygen isotopic exchange with water using the standards USGS34 (δ^15^N of −1.8‰ versus atmospheric N_2_; δ^18^O of −27.9‰ versus Vienna Standard Mean Ocean Water (VSMOW)) and IAEA-3 (δ^15^N of 4.7‰ versus atmospheric N_2_; δ^18^O of 22‰ versus VSMOW) (Böhlke et al., 2003; Frey et al., 2014; McIlvin & Casciotti, 2011).

### Electron balance at oxycline

The balance between the downward flux of electron acceptors (O_2_, NO_3_^-^, NO_2_^-^, and Fe and Mn oxides) and upward flux of electron donors (H_2_S, CH_4_, NH_4_^+^, Mn^2+^, and Fe^2+^) at the oxycline was calculated using the approach described in Żygadłowska et al. (2023). In our calculations, particulate Fe and Mn are assumed to consist of Fe and Mn oxides. The sinking flux of Fe and Mn oxides was assumed to be 0.98 m d^-1^ (Neretin et al., 2003). The vertical eddy diffusivity was estimated from the salinity and temperature profiles. This electron balance was determined to estimate the relative contribution of different oxidants and reductants at the oxycline at the sites with an oxic-anoxic interface in the water column (i.e., Stora Värtan and Skurusundet).

### DNA extractions and 16S rRNA analysis

DNA was extracted with the FastDNA™ SPIN Kit for Soil DNA isolation kit according to the manufacturer’s protocol (MP Biomedicals, Eschwege, Germany). The microbial community composition was analyzed using 16S rRNA gene amplicon sequencing of the V3- V4 region (Illumina MiSeq platform, Macrogen, The Netherlands). The primers used for bacterial 16S rRNA gene amplification were Bac341F (CCTACGGGNGGCWGCAG) (Herlemann et al., 2011) and Bac806R (GGACTACHVGGGTWTCTAAT) (Caporaso et al., 2012). The primers for archaeal 16S rRNA gene amplification were Arch349F (GYGCASCAGKCGMGAAW) (Takai & Horikoshi, 2000) and Arch806R (GGACTACVSGGGTATCTAAT) (Takai & Horikoshi, 2000). To process the sequencing data, Rstudio was used. Sequence quality was assessed with FIGARO and cut according to the FIGARO quality check (Haider et al., 2024). Further trimming and removal of low-quality reads was performed with the DADA2 pipeline (Callahan et al., 2017). Subsequently, amplicon sequence variants (ASVs) were inferred, forward and reverse reads were merged, and chimaeras were removed using DADA2. For taxonomic assignment, the SILVA 16S rRNA gene database (nr99, v138) was used (Quast et al., 2013). The Phyloseq package (McMurdie & Holmes, 2013) was used to cluster ASVs and calculate relative abundances. The data was visualized with the ggplot2 package (Wickham, 2016). Raw reads of the 16S rRNA amplicon sequencing data can be accessed on the European Nucleotide Archive under the accession number PRJNA1126564.

### Metagenome sequencing, assembly, and binning

DNA samples from five water samples (21 m depth for Södra Vaxholmsfjärden; 16 and 19 m for Stora Värtan; 6, 9, and 19 m for Skurusundet) were sequenced (Illumina DNA PCR-Free (450 bp insert) low input kit, Novaseq platform, Macrogen, Amsterdam, The Netherlands). These depths were selected based on water column chemistry. Metagenome assembly and binning were carried out using an adapted version of an in-house bioinformatics pipeline as described in In ‘t Zandt et al. (2019). Briefly FASTQC (v0.11.9; https://www.bioinformatics.babraham.ac.uk/projects/fastqc/) was used for quality assessment. Within the BBMap package (https://sourceforge.net/projects/bbmap/), BBDuk (v37.76) for trimming and filtering of sequencing reads (adapter trimming using the BBDuk-supplied adapters.fa reference with k=23, mink=11, hdist=1, tpe, dbo; quality trimming and filtering using qtrim=lr trimq=15, ftm=5), Tadpole (v39.06) for error correction (mode=correct, k=50), and BBnorm (v37.76) to normalize the coverage of the reads (target=30, min=2). Co-assembly of the remaining reads was conducted by metaSPAdes (v3.5.15) (Nurk et al., 2017) with k-mer sizes 21, 33, 55, 77, 99, 121. Four different algorithms (CONCOCT v1.1.0 (Alneberg et al., 2014), MaxBin2 v2.2.7 (Wu et al., 2016), MetaBAT2 v2.12.1 (Kang et al., 2019), and SemiBin2 v2.0.2 (Pan et al., 2023) were used to bin contigs larger 2,500 bp. Consensus binning was performed using DAS Tool v1.1.1 (Sieber et al., 2018). Final bin quality was assessed with CheckM (v1.2.2) (Sieber et al., 2018), with medium-quality metagenome-assembled genomes (MAGs) defined as >70% complete and <10% contaminated. The taxonomic classification of the bins was obtained with GTDB-Tk (v2.4.0) (Chaumeil et al., 2022). The annotation of the bins, the unbinned fraction, and the contigs <2,500 bp was done with METAscan (Cremers et al., 2022). The raw metagenomic reads can be accessed on the European Nucleotide Archive under the bioproject number PRJNA1126564.

### Gene-centric analysis and bin refinement

The annotated bins, the unbinned fraction, and the contigs <2,500 bp were used to estimate microbial abundances. Contig coverage was calculated using CoverM v0.7.0 (Aroney et al., 2025) and expressed as normalized reads per kilobase per million mapped reads (RPKM). The coverage data were merged with the annotated genes from METAscan, and gene abundance was visualized in R with the ggplot2 package (Wickham, 2016). A genome-centric approach was used to identify the MAGs that have the potential to perform processes within the N-cycle. The MAGs were refined in Anvi’o refine (Anvi’o v8, Eren et al., 2021) and used for further analysis. Selected N-cycle genes of nitrification (*amoA*, *amoB-pmoB*, *amoC-pmoC*, *hao*, *nxrA*), denitrification (*narG*, *napA, nirK*, *nirS*, *norC*, *nosZ),* DNRA *(nrfA*), anammox (*hzsA*), and nitrogen fixation (*nifH*) were extracted from the METAscan output. To visualize the presence of these genes within the MAGs, heatmaps were generated using the heatmap function in R (https://r-graph-gallery.com/heatmap). To determine the relative abundance of MAGs in the samples, CoverM v0.7.0 was used with the setting -relative abundance. Bar plots were generated in R using the ggplot2 or pheatmap packages (https://cran.r-project.org/package=pheatmap) to visualize the results.

## Results and Discussion

### Spatial trends in bottom water redox and sea-air flux of N_2_O

Bottom water euxinia was observed in shallow enclosed areas of the archipelago with relatively little lateral water exchange with the rest of the system (Fig. 1a; Żygadłowska et al., 2024). In the various channels, such as the one connecting the inlet of water from the Norrström River in the Centre of Stockholm with the rest of the archipelago, both oxic and hypoxic bottom waters were found. The continuous surface water N_2_O measurements reveal particularly high concentrations of N_2_O (90 nmol L^-1^) in the waters near the river inlet compared to those in the other parts of the archipelago (10-30 nmol L^-1^; Fig. 1b). Calculated sea-air fluxes of N_2_O show the same pattern with values up to 40 µmol m^-2^ d^-1^ near the inlet and much lower values elsewhere (Fig. 1c). The observed N_2_O concentrations fall within the range previously reported for coastal hypoxic and anoxic systems (4 to 139 nmol L^-1^), which are a known source of N_2_O (Naqvi et al., 2010). The lack of a consistent correlation of high surface water N_2_O concentrations to bottom water hypoxia or euxinia, and the elevated N_2_O concentrations near the river inlet (Fig. 1) indicate multiple controls, which may include nitrification, denitrification, and riverine input, as discussed further below.

### Water column profiles indicate nitrification and denitrification

The three study sites differed strongly in their water column O_2_ and H_2_S profiles (Fig. 2). At Södra Vaxholmsfjärden, the water column was fully oxygenated because of the recent overturn (Żygadłowska et al., 2024). At Stora Värtan, in contrast, an anoxic zone without H_2_S (i.e., a suboxic zone) was present between water depths of 18 and 20 m. Below 20 m, H_2_S was present, with concentrations increasing with depth and reaching a maximum of 44 µmol L^-1^ in the bottom water. At Skurusundet, there was no suboxic zone and the waters became euxinic below 8 m, with H_2_S concentrations at greater depth ranging up to a maximum of 386 µmol L^-1^.

**Figure 2.**
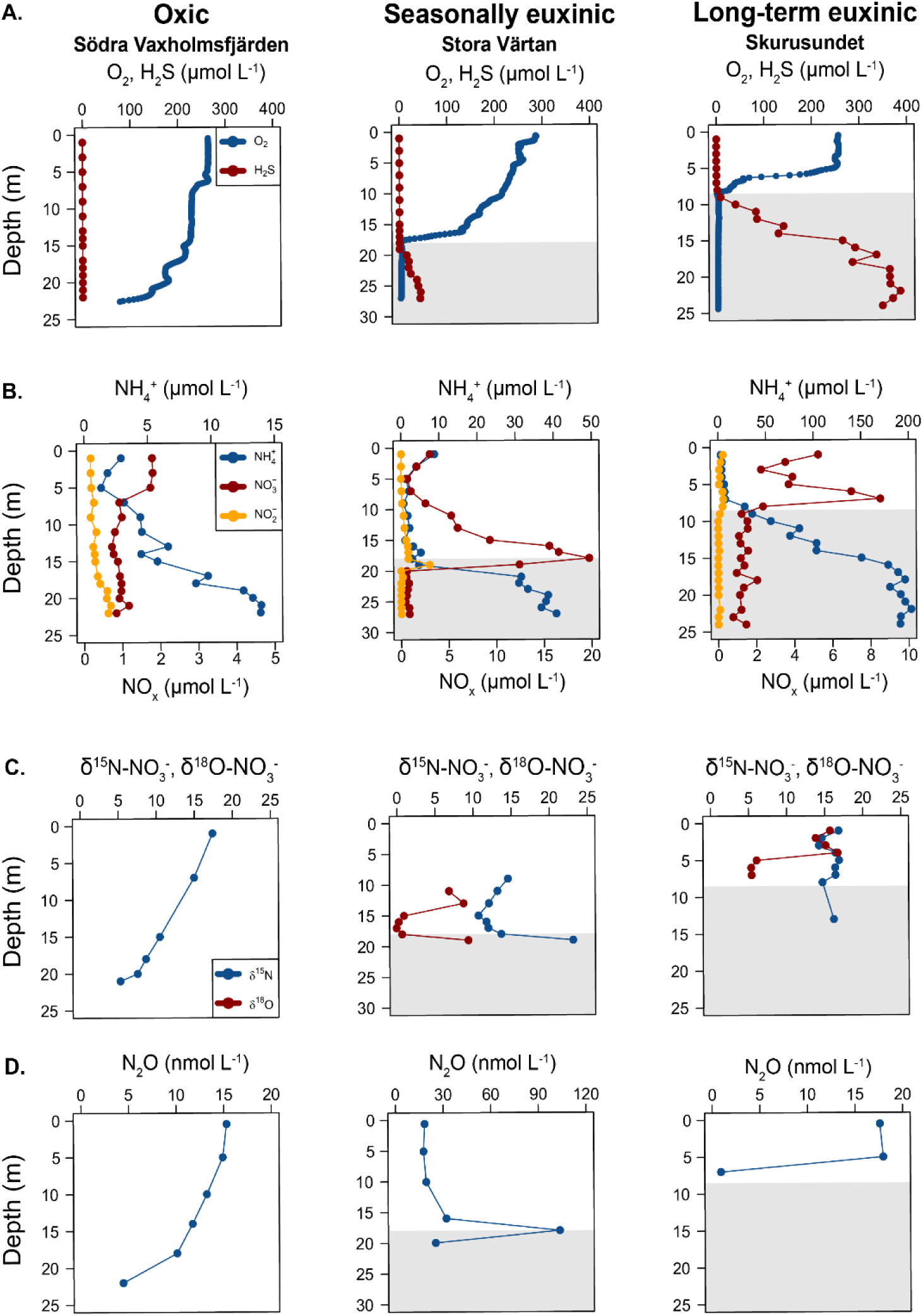
Depth profiles of O_2_ and H_2_S (A.), NH ^+^, NO_2_^-^, and NO_3_^-^ (B.), δ^15^N-NO_3_^-^ and δ^18^O- NO_3_^-^ (C.), and N_2_O (D.) in the water column at Södra Vaxholmsfjärden (oxic; first column), Stora Värtan (seasonally euxinic; second column), and Skurusundet (long-term euxinic; third column). The shading indicates the anoxic part of the water column. N_2_O concentrations are only available for H_2_S-free waters because of methodological constraints. Note the scaling differences of the x-axes showing NH_4_^+^, NO_x_, and N_2_O.

The depth profiles of nutrients were largely in line with the redox conditions. While NH_4_^+^ concentrations at all sites were highest in the bottom water, there was a distinct gradient in concentration from 14 µmol L^-1^ at oxic Södra Vaxholmsfjärden, to 41 µmol L^-1^ at the seasonally euxinic Stora Värtan, and 191 µmol L^-1^ at long-term euxinic Skurusundet (Fig. 2). The upward decrease in NH_4_^+^ concentrations and concomitant increase in NO_3_^-^ and NO_2_^-^ concentrations in the water column suggested that nitrification occurred at all sites.

Nitrification occurred at different depths in the water columns of the three sites. As discussed below, we used the water column solute profiles in combination with the isotopic ^15^N and ^18^O signatures in the NO_3_^-^ pool to deduce the activity of NO_3_^-^ production (nitrification) or removal (denitrification, uptake by phytoplankton) by analysing the depletion and enrichment of the heavy isotopes, respectively (Casciotti, 2016; Lam & Kuypers, 2011). Importantly, lateral inputs can alter these patterns by supplying NO_3_^-^ with another isotopic signature (Naqvi et al., 2010).

At the oxic site Södra Vaxholmsfjärden, counter gradients in NH_4_^+^ and O_2_ suggest a potential for nitrification (Fig. 2a, b). While the upward decrease in NH_4_^+^ might indicate nitrification throughout the water column (Fig. 2b), elevated bottom water NO_2_^-^ concentrations and low values of δ^15^N*-*NO_3_^-^ (Fig. 2b, c) instead suggest that, at this site, most nitrification occurred close to the sediment-water interface. Strikingly, water column NH_4_^+^ concentrations remained elevated throughout the water column (mostly 3 to 14 µmol L^-1^), suggesting inefficient removal (Fig. 2a). The elevated δ^15^N*-*NO_3_^-^ in the surface water might be explained by lateral input, possibly associated with the recent water column mixing (Żygadłowska et al., 2024). Concentrations of N_2_O in the water column ranged from 3 to 15 nmol L^-1^ (Fig. 2d). They decreased near the sediment-water interface, suggesting removal in the sediment, possibly through denitrification, which is the dominant nitrogen removal pathway in the sediment in the region (Van Helmond et al., 2020).

At the seasonally euxinic site Stora Värtan, the upward decrease in water column NH_4_ ^+^ concentrations was accompanied by a peak in NO_3_^-^ with a maximum concentration of 17 µmol L^-1^ at the oxycline (at a depth of 18 m), and a peak in NO_2_^-^ with a maximum concentration of 3 µmol L^-1^ directly below the oxycline (at a depth of 19 m; Fig. 2b). Together with the low values of δ^15^N*-*NO_3_^-^ and δ^18^O-NO_3_^-^ above the oxycline and the high value of δ^15^N*-*NO_3_^-^ at the oxycline (at 18 m; Fig. 2c), this points towards coupled nitrification and denitrification. The sharp maximum in N_2_O observed at the oxycline may result from incomplete nitrification at low O_2_ concentrations or incomplete denitrification (Ward et al., 2008). Taken together, our results suggest that the key nitrogen transformations at this site occur in the suboxic zone between water depths of 18 and 20 m.

At the long-term euxinic site Skurusundet, the upward decrease in NH_4_^+^ concentration was accompanied by relatively small peaks in NO_3_^-^ and NO_2_^-^ above the redoxcline, with maximum concentrations of 8 and 0.3 µmol L^-1^, respectively (Fig. 2b). Together with the low values of δ^18^O-NO_3_^-^ (Fig. 2c), this points to nitrification above the oxycline. The high values of δ^15^N*-*NO_3_^-^ above the oxycline are unexpected, since δ^15^N*-*NO_3_^-^ and δ^18^O-NO_3_^-^ often show similar trends (e.g., Frey et al., 2014) and a rise commonly is interpreted as uptake of NO_3_^-^ (e.g., Frey et al., 2014). The rise in δ^15^N*-*NO_3_^-^ and δ^18^O-NO_3_^-^ in the surface water is likely linked to NO_3_^-^ uptake by phytoplankton. H_2_S was abundantly present in the water column at this site, which interferes with the measurement of N^2^O. Consequently, N_2_O concentrations are only available for the upper part of the water column, where they ranged around 17 nmol L^-1^ (Fig. 2d), falling within the typical range observed in the archipelago (Fig. 1).

In summary, the water column chemistry shows that nitrification occurs in the water column at all three sites. While NH_4_^+^ removal appears relatively inefficient at the oxic Södra Vaxholmsfjärden site, the peaks in NO^3-^ concentrations and the isotopic signatures of NO_3_^-^ in the waters near the oxycline at the other two stations suggest coupled nitrification and denitrification. When taking the height of the NO_3_^-^ peak as an approximation for nitrification activity, the seasonally euxinic Stora Värtan site, which has a suboxic zone, appears to be the most active regarding N cycle processes.

### 16S rRNA gene and metagenomic-based evidence for microbial nitrogen cycle processes

Amplicon sequencing profiles of 16S rRNA genes provided insight into the distribution and abundance of microorganisms contributing to N cycle processes at the investigated sites in the Stockholm Archipelago (Fig. 3a). At the oxic site (Södra Vaxholmfjärden), the relative abundance of nitrifier-associated 16S rRNA genes increased with water depth, which is consistent with the water column chemistry-inferred nitrification in the bottom water. This trend was not observed for *Nitrosopumilis*, but, based on metagenomic read mapping, the abundance of archaea is too low to deduce depth trends accurately. The 16S rRNA gene profiles for the other two sites show that nitrifiers had the highest relative abundance at the oxycline. This is particularly apparent for seasonally euxinic site Stora Värtan, where AOB, AOA, and NOB 16S rRNA gene abundances peak between 15 and 21 meters depth (Fig. 3a). Concurrent with the depth at which H_2_S concentrations start to increase, their abundance decreases below 21 meters. Contrastingly, the 16S rRNA gene-based relative abundances of *Sulfurimonas* increase between 16 and 20 meters and remain constant in the sulfidic bottom waters (Fig. 3b). *Sulfurimonas*- related bacteria can grow on diverse sulfur compounds and can be key players in marine denitrification (Shao et al., 2011; Takai et al., 2006), and, accordingly, the genome of the *Sulfuriomonas* species found here possessed denitrification genes (see below). Furthermore, as it were the most abundant MAGs with denitrifying potential, it can be used as a proxy for denitrification. At the permanently euxinic site Skurusundet, the 16S rRNA gene-based relative abundances of nitrifiers showed a small increase between depths of 6 and 8 meters, directly above the oxycline (Fig. 2 and 3a). However, they are present at only low levels in the anoxic and sulfidic zone below 8 meters. In the euxinic water column, *Sulfurimonas*-related species again constituted a large portion of the microbial population (Fig. 3b and S1).

**Figure 3.**
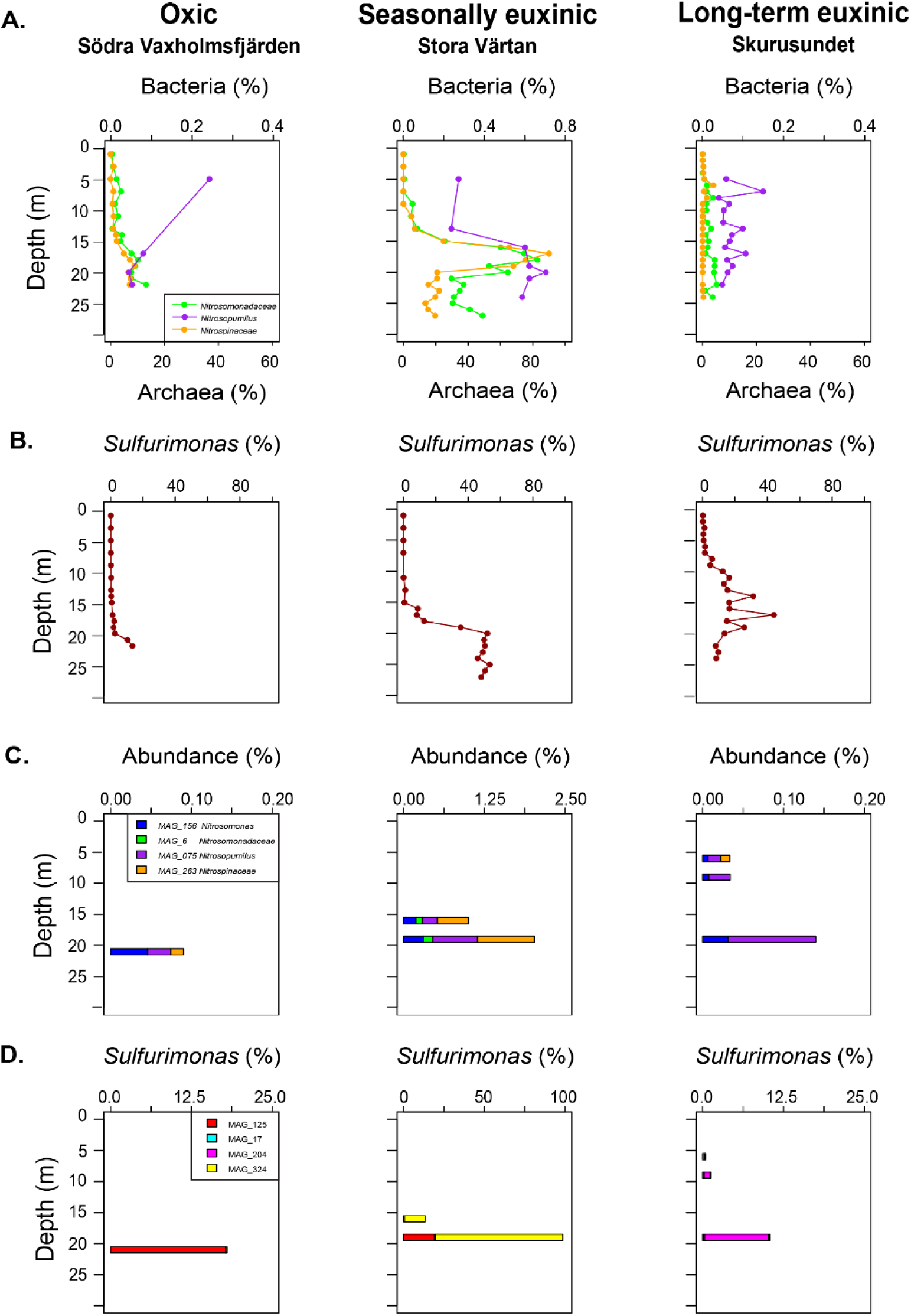
Depth profiles of (a) 16S rRNA gene-based relative abundances of nitrifying bacteria and archaea (Nitrosomonadaceae, Nitrosopumilus and Nitrospinaceae,) and Sulfurimonas as a proxy for denitrification throughout the water column and (b) metagenome-assembled genome-based abundances of nitrifiers and Sulfurimonas at Södra Vaxholmsfjärden at 21 m, Stora Värtan at 16 m and 19 m, and Skurusundet at 6 m, 9 m, and 19 m depth.

In addition to 16S rRNA gene amplicons, we obtained metagenomic data for water column samples from Södra Vaxholmsfjärden at 21 meters, Stora Värtan at 16 and 19 meters, and Skurusundet at 6, 9, and 19 meters depth. In total, we obtained four MAGs belonging to nitrifiers. Two MAGs belonged to *Nitrosomonas*-like AOB affiliated with the *Nitrosomonadaceae* family, one MAG to a *Nitrosopumilus*-like AOA, and one NOB within the *Nitrospinaceae*. These nitrifiers had the highest abundances at Stora Värtan compared to the other sites, with the *Nitrospinaceae* MAG reaching the highest abundance of 0.48% and 0.88% at 16 m and 19 m (Fig. 3c), respectively. Sulfide-oxidizing nitrate-respiring *Sulfurimonas* species were abundant in the water columns of all three sites, and four good-quality MAGs were retrieved. Interestingly, at each station, distinct *Sulfurimonas* MAGs were dominant (Fig. 3d).

We furthermore investigated all retrieved MAGs for their N-cycling potential (Fig. 4). The *Nitrosomonadaceae* AOB MAG_6, which was classified to the family level only, contained the *amoCAB*, *hao*, and, additionally, *nirK* genes. In contrast, MAG_156 was classified as *Nitrosomonas* but lacked *amo* and *hao* genes, and we could not resolve the absence of nitrification genes by manual MAG refinement. It did, however, contain *nirK* and *norC*. The *Nitrosopumilus* MAG_075 included, after refinement, the *amoA, amoC*, and nirK genes. Most *Sulfurimonas* MAGs, except for *Sulfurimonas* MAG_324, only encoded a partial denitrification pathway. In addition, several taxonomically diverse MAGs contained denitrification genes (Fig. 4). The *Desulfuromusa* MAG_019 had the genomic potential for DNRA as indicated by *nrfA*, and five MAGs showed potential for nitrogen fixation based on the presence of *nifH* (Fig. 4).

**Figure 4.**
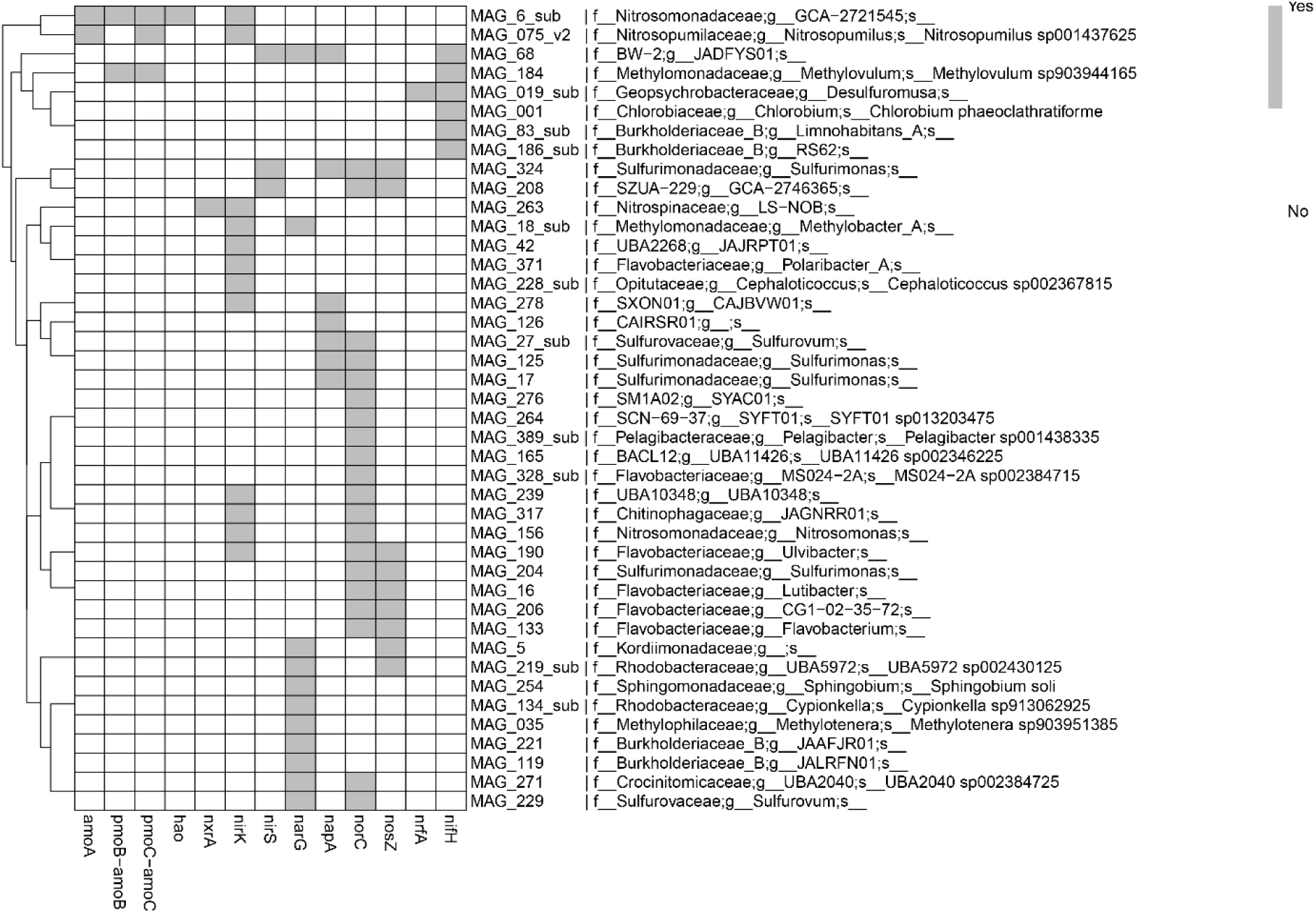
Presence of N-cycling genes in the obtained MAGs. The presence of genes is indicated in grey and their absence in white. The MAGs are clustered according to gene content.

We also investigated the distribution of N cycle processes using a gene-centric approach to analyse the relative abundance of the diagnostic genes for nitrification, denitrification, DNRA, and N-fixation (Fig. 5), as the marker gene for anammox (*hzsA*) bacteria was not detected at any site. Congruent with the MAG-based analysis, the highest relative abundance of nitrification genes was found at Stora Värtan, where the denitrifier marker genes *nirS*, *norC*, and *nosZ* also had high RKPM values. Södra Vaxholmsfjärden shows the lowest RKPM values for N cycle genes, with *norC* being the most abundant. At Skurusundet, the RKPM values are slightly lower than at Stora Värtan, and the sulfidic waters below the oxycline appear to contain more genes involved in dissimilatory nitrate reduction than the other depths analysed at this site. Interestingly, the nitrogen fixation marker gene *nifH* has the highest RKPM value of all samples in this ammonium-rich zone at Skurusundet.

**Figure 5.**
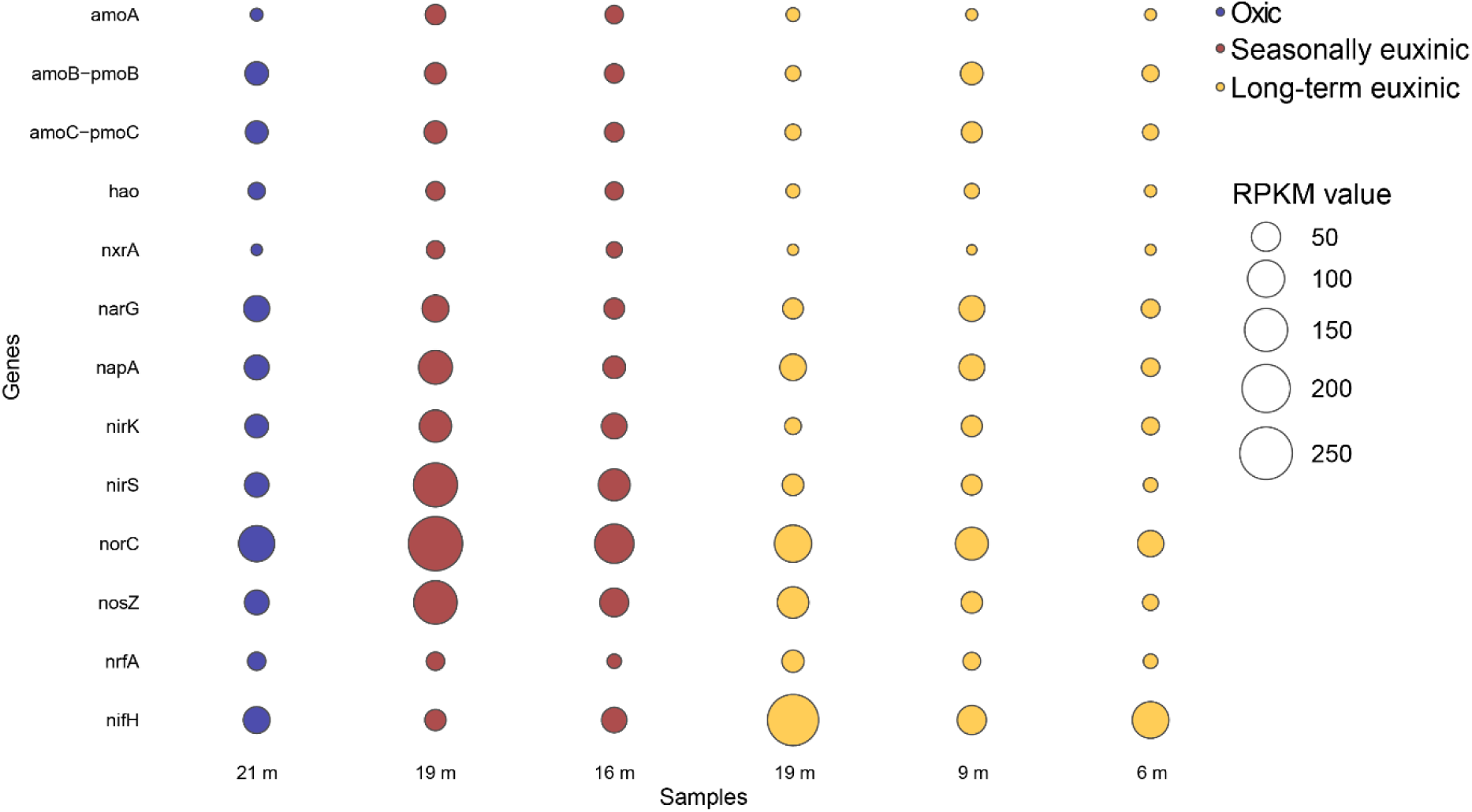
Relative abundance of N cycle genes at Södra Vaxholmsfjärden (VAX) at 21 m depth in blue, Stora Vartän (VAR) at 16 m and 19 m depth in red, and Skurusundet (SKU) at 6 m, 9 m, and 19 m depth in yellow. The bubble size represents the normalized gene abundance in reads per kilobase per million mapped reads (RPKM).

### Stratification and euxinia as key controls on nitrification and denitrification

All three sites investigated in the Stockholm archipelago showed distinct differences in water column N cycling, as reflected in the nutrient profiles, nitrogen isotope distributions, and microbial community composition. Our results suggest that these differences are primarily controlled by the degree and duration of water stratification and associated changes in bottom water redox conditions.

At Södra Vaxholmsfjärden, where the recent overturn led to oxygenation of the water column (Żygadłowska et al., 2024), both the relatively low NO_3_^-^ concentrations and low relative abundances of nitrifiers indicated limited nitrification. An explanation could be that the nitrifiers in this brackish system are adapted to low O_2_ concentrations and hence are inhibited by high O_2_ levels in the water column (Goreau et al., 1980; Hulth et al., 2005) In addition, the recent physical mixing of the water column may have hindered nitrification through biomass dilution, as shown for waters of a fjord-like coastal system in Canada (Haas et al., 2021).

The difference in duration of stratification at Stora Värtan and Skurusundet had major consequences for the water column chemistry. This is illustrated with an electron balance for the oxycline at both sites in which the fluxes of electron donors and acceptors are compared (Supplementary Table S1). O_2_ accounted for the majority of the downward oxidant flux at Stora Värtan and Skurusundet (74% and 92%, respectively), with an additional contribution from NO_3_^-^ (19.6% and 6.8%, respectively). NH_4_^+^ and H_2_S together accounted for ∼75% and 86% of the total reductant flux at Stora Värtan and Skurusundet, respectively, and organic matter in the water column might have contributed to the redox balance. Still, the results point towards an excess of electron donors over electron acceptors at both sites, indicating a shallowing of the oxycline and increasingly sulfidic waters upon increased stratification duration.

The seasonally euxinic site Stora Värtan is characterized by a much higher abundance of nitrifiers near the oxycline than the long-term euxinic Skursundet (Fig. 3). This is in line with the more distinct and higher NO_3_^-^ and NO_2_^-^ peaks near the oxycline (Fig. 2) and a greater downward NO_3_^-^ flux into the anoxic zone at the former site. This suggests higher rates of nitrification at Stora Värtan compared to Skursundet. This difference is likely the result of the more abundant presence of H_2_S at Skursundet. H_2_S is known to inhibit bacterial nitrification (Joye & Hollibaugh, 1995; Lam et al., 2007), and we indeed find a distinct decrease in the relative abundance of *Nitrosomonadaceae* with increased H_2_S availability (Fig. 3). The inhibitory effect of H_2_S is known to be less pronounced for AOAs. For example, in the Baltic Sea, where H_2_S concentrations are generally high near the oxycline, *Nitrosopumilus* is often the sole ammonia oxidizer present (Berg et al., 2015; Jäntti et al., 2018; Labrenz et al., 2010).

Although archaea had a low abundance in the water column, we detected *Nitrosopumilus* at all three sites, also below the oxycline and in sulfidic waters (Fig. 3). A recent study showed that the AOA *Nitrosopumilus maritimus* may produce O_2_ via NO dismutation, which is subsequently used for ammonia oxidation and may explain its presence in anoxic water (Kraft et al., 2022). However, more research is needed to understand the nitrifiers’ metabolism below the oxycline in coastal waters. Regardless, most nitrification in coastal systems is expected to take place at or near the oxycline where NH_4_^+^ and O_2_ meet (e.g., Jantti et al., 2018), and, as highlighted in our study, this process is hindered by water column euxinia.

In contrast, H_2_S stimulated denitrification at the investigated sites. This is most evident from the NO_3_^-^ and NO_2_^-^ peaks around the oxycline at Stora Värtan and Skurusundet (Fig. 2) and the high relative abundance of *Sulfurimonas* species, which seems to be a good proxy for denitrifying activity since these bacteria possess the genetic potential for denitrification (Fig. 3). Previous studies have shown that *Sulfurimonas* is also the dominant denitrifier at the oxycline in the Central Baltic Sea (Bruckner et al., 2012; Glaubitz et al., 2008; Grote et al., 2007). The dominant denitrifier at Stora Värtan, *Sulfurimonas* MAG_4, has the potential for full denitrification, making it striking that there is a peak in N_2_O at the oxycline at this site. N_2_O production could result from copper limitation, preventing N_2_O reductase from functioning (Ward et al., 2008), or oxygen-limited nitrification (Kozlowski et al., 2016).

In addition to denitrification, there is also potential for DNRA, as indicated by the presence of *nrfA* genes in the bottom waters of all sites. Although potential rates for DNRA have been measured in the Baltic Sea, this process is considered to be of minor importance relative to denitrification (Bonaglia et al., 2016). We found no indications for anammox in the 16S rRNA gene or the metagenomic data at any site. Since anammox bacteria are slow-growing and the water column at the oxic and hypoxic study sites is not permanently stratified, they may be unable to establish. Furthermore, anammox bacteria are known to be very sensitive to H_2_S (Russ et al., 2014). A study in the Black Sea showed, for example, that anammox activity is inhibited at H_2_S concentrations as low as 1.5 μmol L^-1^ (Jensen et al., 2008). In the Central Baltic, anammox has been shown to occur following an inflow of oxygenated North Sea water that allows the formation of a suboxic zone (Hannig et al., 2007). Recent work found anammox to occur at low rates during periods of stagnation if a suboxic zone is present (Bonaglia et al., 2016).

Interestingly, a high abundance of *nifH* was observed at Skurusundet (Fig. 5). The *Chlorobium phaeclathratiforme* MAG_001, which contains *nifH*, is highly abundant according to this site’s 16S rRNA sequencing results. Intriguingly, this is a photolithotrophic green sulfur bacterium that is capable of photosynthesis at extremely low light intensities (Raven et al., 2000). *Chlorobium* abundances continue to increase with water depth and remain abundant in the bottom water. Although *nifH* is present in high abundance, the high NH_4_^+^ in the bottom water makes it unlikely that the gene is expressed, but further studies are needed to investigate nitrogenase activity and the role of diazotrophs in this system.

## Conclusions

In this study, we combined chemical and microbial analyses to assess nitrogen dynamics in a eutrophic coastal system at three different sites with contrasting redox conditions. Nitrification was promoted at the redox interface and appeared to be inhibited in a fully oxic water column. Denitrification actively contributed to nitrogen removal under anoxic and euxinic conditions, whereas anammox bacteria were absent at all three sites. The activity and abundance of nitrogen-cycling microorganisms, especially nitrifying and denitrifying bacteria, varied across the three sites, and no nitrifiers were detected in the euxinic site. Long-term euxinia caused a decrease in nitrifier abundance despite the presence of NH_4_^+^, and consequently promoted NH_4_^+^ accumulation over removal. Prolonged euxinia thus sustains eutrophication and deoxygenation of coastal systems.

## Acknowledgements

We thank the captain and crew of the R/V Electra for their support during the sampling campaigns. We are also grateful to J. Visser for analytical assistance.

## Financial support

This research was financially supported by ERC Synergy grant Marix (854088) and by the Swedish Research Council (VR) grants 2021-04641, 2018-04350, and 2022-04081. This work was carried out under the Netherlands Earth System Science Center program (NESCC 24002001), supported by the Ministry of Education, Culture, and Science (OCW).

## Notes

### Competing Interest Statement

The authors have declared no competing interest.

