## Supplementary Information for "Long-term euxinia hinders microbial ammonium removal in brackish coastal waters"

#### Flux determination for N_2_O

The diffusive flux of N­_2_O from the water column to the atmosphere was determined as:

F = k (C_W_ – C_O_),	(1)

where F is the sea-air flux of N_2_O (µmol m^-2^ d^-1^), k is the gas exchange coefficient of N_2_O (m d^-1^), C_W_ is the concentration of N_2_O in seawater (µmol L^-1^), and C_O_ is the concentration of N_2_O in equilibrium with the atmosphere (µmol L^-1^) (Garbe et al., 2014). The concentrations of N_2_O in seawater and air were obtained following the approach of Żygadłowska et al. (2024):

C_W_ = xN_2_O_sw_ * ß * P	(2a)

C_O_ = xN_2_O_atm_ * ß * P,	(2b)

where xN_2_O_sw_ is the concentration of N_2_O at 1 m depth obtained from the continuous surface water measurements along the transect, xN_2_O_atm_ is the concentration of N_2_O in the atmosphere at the time of sampling obtained from the Global Monitoring Laboratory (<https://gml.noaa.gov/dv/data.html>) using the nearest station (i.e. Mace Head), *ß* is the Bunsen solubility (dimensionless) and P is the atmospheric pressure (bar). The Bunsen solubility is calculated using the coefficients and equation from Weiss and Price (1980):

*ln ß* = A_1_ + A_2_ $\left( \frac{100}{T} \right)$+ A_3_ ln$\left( \frac{100}{T} \right)$+ S [ B_1_ + B_2_ + $\left( \frac{100}{T} \right)$+ B_3_ $\left( \frac{100}{T} \right)$^2^],	(3)

where A_1_, A_2_, A_3_, B_1_, B_2_, B_3_ are constants (mol L^-1^ atm^-1^), T is the absolute temperature (K), and S is the salinity in parts per thousand (‰).

The gas exchange coefficient (k) was calculated using Wanninkhof (2014):

k = 0.251 * U ^2^ * (Sc/660)^-0.5^,	(4)

where U is wind speed at 10 m in height (cm h^-1^) and Sc is the Schmidt number (dimensionless). The windspeed at the time of sampling was obtained from Swedish Meteorological and Hydrological Institute (SMHI; https://www.smhi.se/data/meteorologi/). The Schmidt number was calculated at in situ conditions by dividing the kinematic viscosity of seawater (*v*) by the diffusion coefficient (*D*) of N_2_O in seawater (Morgan et al., 2019). The kinematic viscosity of seawater at in situ conditions was taken from Pilson (2013). The diffusion coefficient is derived using the method described in Bange et al., (2001):

D_N2O_ = 3.16*10^-6^ exp (-18.370/RT),	(5)

where R is the universal gas constant and T is the absolute temperature (K).

### Supplementary Figures and Tables

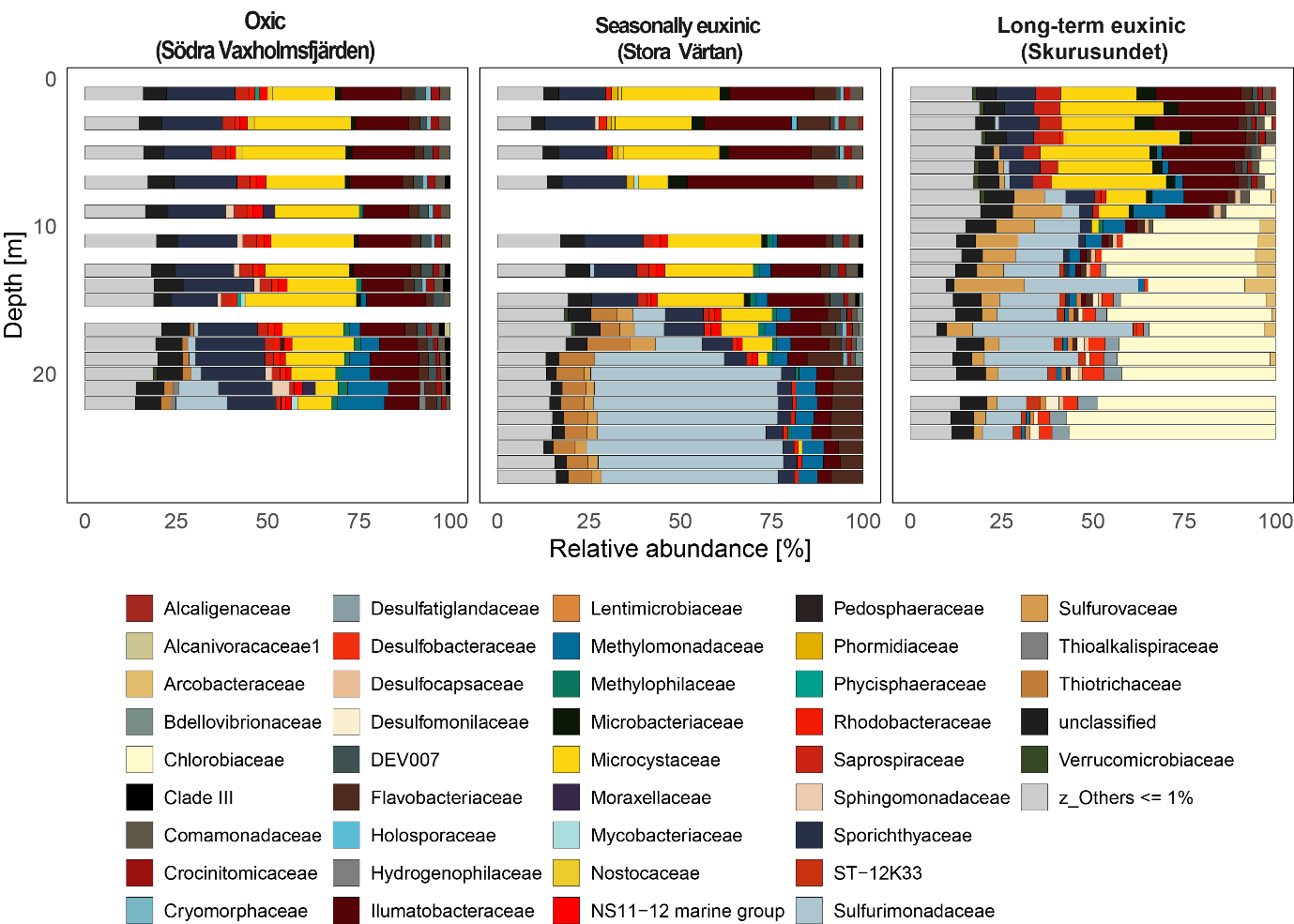

**Figure S1.** 16S rRNA gene amplicon sequencing results for bacterial genera with more than 1% relative abundance. Genera lower than 1% are summarized as “Others”.

**Table S1.** Balance of electron acceptors (oxidants) and electron donors (reductants) at the oxycline at Stora Värtan and Skurusundet. Negative and positive values refer to downward and upward fluxes, respectively.

|  | | **Electron acceptors** | | | | | | **Electron donors** | | | |
| --- | --- | --- | --- | --- | --- | --- | --- | --- | --- | --- | --- |
|  |  | **O_2_** | **Mn(IV)** | **Fe(III)** | **NO_3_^-^** | **NO_2_^-^** | **H_2_S** | **CH_4_** | **NH_4_^+^** | **Mn(II)** | **Fe(II)** |
|  | Transferable  electrons | 4 | 2 | 1 | 5 | 3 | 8 | 8 | 8 | 2 | 1 |
| ***Störa Värtan*** | Flux  (mmol m^-2^ d^-1^) | -11.67 | -0.67 | -0.37 | -2.48 | -0.85 | 5.04 | 3.13 | 5.75 | 2.08 | 0.19 |
|  | Electron flux  (mmol e^-^ m^-2^ d^-1^) | -46.67 | -1.35 | -0.37 | -12.39 | -2.56 | 40.30 | 25.06 | 45.97 | 4.15 | 0.19 |
|  | Electron contribution  (%) | 73.7 | 2.1 | 0.6 | 19.6 | 4.0 | 34.8 | 21.7 | 39.7 | 3.6 | 0.2 |
|  | **Electron balance** | **-63.33** | | | | | | **115.68** | | | |
|  |  | **52.35** | | | | | | | | | |
| ***Skurusundet*** | Flux  (mmol m^-2^ d^-1^) | -30.56 | -0.58 | -0.32 | -1.80 | -0.02 | 11.12 | 3.04 | 7.43 | 0.24 | 0.22 |
|  | Electron flux  (mmol e^-^ m^-2^ d^-1^) | -122.26 | -1.16 | -0.32 | -9.02 | -0.07 | 88.95 | 24.34 | 59.47 | 0.48 | 0.22 |
|  | Electron contribution  (%) | 92.0 | 0.9 | 0.2 | 6.8 | 0.1 | 51.3 | 14.0 | 34.3 | 0.3 | 0.1 |
|  | **Electron balance** | **-132.83** | | | | | | **173.45** | | | |
|  |  | **40.62** | | | | | | | | | |
